# Cerebellar anodal tDCS increases implicit visuomotor remapping when strategic re-aiming is suppressed

**DOI:** 10.1101/091397

**Authors:** Li-Ann Leow, Welber Marinovic, Stephan Riek, Timothy J Carroll

## Abstract

The cerebellum is known to be critically involved in sensorimotor adaptation. Changes in cerebellar function alter behaviour when compensating for sensorimotor perturbations, as shown by non-invasive stimulation of the cerebellum and studies involving patients with cerebellar degeneration. It is known, 24 however, that behavioural responses to sensorimotor perturbations reflect both explicit processes (such as volitional aiming to one side of a target to counteract a rotation of visual feedback) and implicit, error-driven updating of sensorimotor maps. The contribution of the cerebellum to these explicit and implicit processes remains unclear. Here, we examined the role of the cerebellum in sensorimotor adaptation to a 30° rotation of visual feedback of hand position during target-reaching, when the capacity to use explicit processes was manipulated by controlling movement preparation times. Explicit re-aiming was suppressed in one condition by requiring subjects to initiate their movements within 300ms of target presentation, and permitted in another condition by requiring subjects to wait approximately 1050ms after target presentation before movement initiation. Similar to previous work, applying anodal transcranial direct current stimulation (tDCS; 1.5mA) to the right cerebellum during adaptation resulted in faster compensation for errors imposed by the rotation. After exposure to the rotation, we evaluated implicit remapping in no-feedback trials after providing participants with explicit knowledge that the rotation had been removed. Crucially, movements were more adapted in these no-feedback trials following cerebellar anodal tDCS than after sham stimulation in both long and short preparation groups. This suggests that cerebellar anodal tDCS increased implicit remapping during sensorimotor adaptation irrespective of preparation time constraints. This work shows that the cerebellum is critical in the formation of new visuomotor maps that correct perturbations in sensory feedback, both when explicit processes are suppressed and when allowed during sensorimotor adaptation.

## Cerebellar anodal tDCS increases implicit visuomotor remapping when strategic re-aiming is suppressed

The cerebellum has long been known to play a crucial role in predicting the sensory consequences of motor commands [1]; a process that appears necessary both for rapid online responses to unexpected events, and for trial-by-trial compensation of systematic sensorimotor disturbances (for recent reviews, 48 see [2, 3]). When a perturbation of sensory feedback (e.g., a rotation in visual feedback of a movement trajectory, or a force field that pushes the moving hand away from its intended direction) evokes a mismatch between the predicted sensory outcomes and the actual sensory outcomes, the internal mapping between motor commands and resulting changes in sensory state is thought to be updated, such that the prediction error is minimized in subsequent movements. The likely involvement of the cerebellum in this process is supported by a large body of computational, neurophysiological and neuropsychological work. For example, patients with selective degeneration of the cerebellum show substantially impaired capacity to correct for various different types of perturbations, including velocity-dependent force-fields [4-6], 56 translated feedback of the entire visual field [7-9], rotated visual feedback of hand movement trajectories [10, 11], or adaptation of walking to differing speeds imposed on the left and right legs in split-belt treadmill adaptation [12, 13].

People compensate for systematic sensorimotor perturbations, either by using an explicit strategy to alter their movement characteristics (e.g., explicitly aiming in a different direction from the target), or through implicit learning of new sensory-motor mappings [14, 15]. It has been proposed that the cerebellum is not required to strategically modify movements, as patients with cerebellar degeneration can use re-aiming strategies when explicitly instructed how to do so [16]. Although this suggests that the cerebellum is not crucial in implementing strategic compensations to perturbations, the observation that patients typically do not spontaneously develop such strategies implies that the cerebellum may play a role in identifying or formulating strategies [16]. Importantly, it has been shown that the initial rapid rate of error reduction in sensorimotor adaptation tasks is dominated by the explicit component of sensorimotor adaptation [14, 17]. Several studies have shown that increasing the excitability of the cerebellum via non-invasive brain stimulation increases the rate of the initial rapid error reduction in sensorimotor adaptation tasks [18-22], 70 raising the possibility that faster error compensation with cerebellar anodal tDCS occurs in part by upregulating explicit processes.

No previous studies have assessed whether cerebellar anodal tDCS affects the rate or extent of sensorimotor adaptation when explicit compensatory processes are suppressed. One way to dissociate implicit and explicit mechanisms during adaptation is to reduce the amount of time available to prepare movement [23-25], because employing explicit re-aiming strategies is difficult under time pressure, and longer movement preparation time is associated with the employment of explicit re-aiming strategies [26, 27]. Here, we examined whether cerebellar anodal tDCS affected adaptation to a 30° rotation of visual feedback of the movement trajectory when explicit processes were suppressed by enforcing short preparation times [24]. Crucially, we quantified the extent to which participants acquired new sensorimotor maps in trials where participants reached towards targets without visual feedback of movements, and with knowledge that the rotation had been removed. Cerebellar anodal tDCS increased the extent of *implicit* remapping resulting from exposure to the perturbation in both preparation time conditions. This suggests that the cerebellum contributes to implicit sensorimotor remapping regardless of whether explicit strategy use is suppressed or allowed during learning.

## Method

### Participants

Seventy-two right-handed individuals (mean age= 22.2, years SD=2.85) completed the study. We decided to collect a minimum of 14 datasets for each condition a-priori, based on our previous study which showed reliable effects of manipulating preparation time with 14 participants in short and long preparation time conditions [25]. For each participant group, half of the participants were randomly assigned to clockwise and counter-clockwise conditions. Data from four participants were excluded from the analysis: due to experimenter error for three participants (two received incorrect task instructions, one did not complete a baseline phase), and due to voluntary dropout in one participant. No other datasets were removed from the analyses. The final sample sizes for each experimental condition were as follows: Cerebellar Anodal Short Preparation time (n=15, 7 counterclockwise, 8 clockwise), Cerebellar Sham Short Preparation Time (n=14, 7 counterclockwise, 7 clockwise), Cerebellar Anodal Long Preparation Time (n=21, 10 counterclockwise, 11 clockwise), Cerebellar Sham Long Preparation Time (n=20, 9 counterclockwise, 9 clockwise). All participants were naïve to visuomotor rotation and force-field adaptation tasks. Participants were reimbursed with course credits or with monetary reimbursement of $10 per hour of participation. The experiments were approved by the Ethics Committee of the University of Queensland and are in accordance with The Declaration of Helsinki.

### tDCS

Prior to behavioral testing, the scalp area overlying the right cerebellum was localized using the international electroencephalographic 10 – 20 system. For all groups, the anodal electrode was placed over the scalp area estimated to overly the right cerebellar cortex (3 cm lateral to the inion), and the reference electrode was positioned on the skin area overlying the right buccinator muscle[28]. This method of localizing the right cerebellum has been found to be appropriate for tDCS of the right cerebellum. 4.5 x 4.5 cm carbon-rubber electrodes were encased in saline soaked sponge pads (4.5 cm x 6 cm, Soterix Medical Inc. EasyPAD), and secured using Velcro straps, and stimulation was generated with a Soterix (Soterix Medical Inc., NY) (current density of approximately 0.08 mA/cm2). The current was gradually ramped up to 1.5 mA over 30 s starting from the last 10 baseline trials prior to the adaptation block, before the initial block of adaptation trials. The stimulation lasted the entire adaptation block, or a maximum of 40 minutes, whichever came sooner, and then was gradually ramped down over 30s. For the sham tDCS conditions, the current was ramped down over a 30 s period immediately after achieving the maximum of 1.5mA.

### Apparatus

Participants completed the task using the VBOT planar robotic manipulandum, a custom-built planar robotic interface with a low-mass, two-link carbon fibre arm which measures position with optical encoders sampled at 1,000 Hz. For more details of the experimental setup, see Howard, Ingram (29). Participants made centre-out horizontal reaching movements by moving the handle of the manipulandum to move an on-screen circular cursor (radius 0.25cm) from a start circle (radius 0.5cm) to a target circle (radius 0.5cm), projected on a computer monitor (ASUS, VG278H, Taiwan) running at 60Hz mounted above the vBOT via a mirror in a darkened room. Participants observed the monitor via its reflection onto a horizontal mirror which prevented direct vision of their arm, and gave the illusion that the cursor and targets were located in the plane of hand motion. Participants were seated on a chair height-adjusted to allow optimal viewing of the screen for the duration of the experiment. The right forearm was supported by an air-sled which rested on a glass table. Compressed air was forced out of small holes in the air-sled runners, which allowed low friction in the plane of movement. Targets appeared randomly in one of eight locations (0°, 45°, 90°, 135°, 180°, 225°, 270° and 315° relative to the start circle located centrally on-screen). The distance from the center of the start circle to the center of the targets was 9cm.

### General Trial Structure

Participants were instructed that their goal was to move the cursor (radius 0.25cm) as accurately as possible from the start circle (radius 0.5 cm) to the target circle (radius 0.5cm). Participants were instructed not to stop on the target, but to slice through the target. Across all conditions, a sequence of three tones spaced 500 ms apart were presented at a clearly audible volume via external speakers. Participants were instructed to time the onset of their movements with the onset of the third tone. This timed-response paradigm has previously been shown to be effective in encouraging adherence to stringent response time requirements [30-33]). Movement initiation was defined online as when hand speed exceeded 2cm/s. Targets appeared at 1000ms (long preparation time condition) or 250 ms minus a display latency (27.6 ± 1.8 ms), prior to the third tone. Thus target direction information became available 972.4 or 222.4ms before the desired initiation time. When movements were initiated 50 ms later than the third tone, the trial was aborted: the screen was blanked and a “Too Late” on-screen error signal appeared. Similarly, when movements were initiated more than 100 ms before the desired initiation time, the trial was aborted: the screen was blanked and a “Too Soon” on-screen error signal appeared. No visual feedback about movements was available when trials were aborted. Thus, all movements recorded and analysed were made according to the following “hard cut-off” times: within 1022.4 ms after target presentation for the long preparation time condition, and within 272.4 ms after target presentation for the short preparation time condition.

Participants in all conditions first completed a **baseline** pre-rotation block of 6 cycles (48 trials) with veridical feedback of their movement trajectories via on-screen cursor position to familiarize them with the task. The baseline block was followed by an adaptation block (60 cycles, i.e., 480 trials) with either a 30° clockwise or counterclockwise rotation of visual feedback relative to the center of the start circle. The **adaptation** block was followed by a **no-feedback** block of 6 cycles (i.e., 48 trials), where visual feedback of cursor position was hidden immediately after the cursor left the start circle. Crucially, before commencing this block, participants were explicitly instructed that there was no longer any disturbance of visual feedback, and that they should aim straight towards the target [14, 34]. The residual learning that remained after removing the influence of explicit learning is therefore assumed to be implicit in nature-this no-feedback block is therefore thought to assay implicit acquisition of new sensorimotor maps (thereafter termed **implicit remapping)**. Finally, participants completed a **washout** block of 6 cycles (48 trials) where unrotated visual feedback of cursor position was available to enable participants to return movements to the unadapted state. The same preparation time constraints were maintained throughout the entire experiment for each group.

### Data analysis

Movement onset time was taken as the time at which hand speed first exceeded 2 cm/s. Movement direction was quantified at 20 percent of the movement distance. This procedure ensured that movement direction was quantified at less than 200ms into the movement, at which time the size of online corrections in hand position is small [35].

Intrinsic biases in reaching direction can affect adaptation behaviour [36-38]. Intrinsic biases were estimated by averaging movements from the last baseline cycle that were within 90° of the target (i.e., 45° clockwise or counterclockwise of the target). Then, this (estimated) bias was subtracted from movement direction for each trial. Trials were averaged in cycles of 8 trials (one cycle for each of the 8 target angles) for analysis. In the adaptation, no-feedback, and washout blocks, data for participants who experienced counterclockwise rotations (-30°) were sign-transformed and collapsed for analysis with data for participants who experienced clockwise (°30°) rotations. We did not apply any outlier removal procedure for the adaptation phase, the no-feedback phase, and the washout phase.

For the adaptation phase, we defined adaptation into an early phase and a late phase by splitting the 60- cycle adaptation block into two phases: the early phase (Cycles 1-30), and the late phase (Cycles 31-60). Separate ANOVAs with between-subjects factors Stimulation (cerebellar anodal tDCS, cerebellar sham tDCS) and Preparation Time (short preparation time, long preparation time) and within-subjects factors Cycle were run for the early and the late phase. To evaluate implicit remapping after attaining explicit knowledge that the rotation had been removed, we ran Stimulation (cerebellar anodal tDCS, cerebellar sham tDCS) x Preparation Time (short preparation time, long preparation time) x Cycle ANOVAs, and a separate stimulation (Sham, Stim) x Preparation Time (Short, Long) ANOVA on the No Feedback block. For these mixed-ANOVAs, rotation direction was included as a variable of no interest in our ANOVAs, as rotation direction was not part of our hypotheses, and multi-way mixed ANOVAs with a large number of factors have an increased likelihood of generating spurious results (Cramer et al. 2016).

In addition, to examine the rate of adaptation without the possible confound of intrinsic bias in movement direction, we also fit cycle-averaged movement directions for each dataset to a single-rate exponential function [39], as follows:

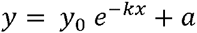

where *y* is the movement direction, *x* is the trial number, *k* is the rate constant that indicates the rate with which movement direction changes, *a* is the movement direction at which performance reaches asymptote, and *y_0_ ° a* is the hypothetical *y* value when *x* is zero.

We also examined the rate of de-adaptation in the washout block by fitting cycle-averaged movement directions for the washout block to a straight line, as follows:

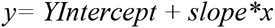

where y is the movement direction, x is the trial number, slope is the rate constant that indicates the rate with which movement direction changes, and YIntercept is the hypothetical y value when x is zero.

GraphPad 7.0 least squares non-linear regression was used to fit data to both functions. Non-linear regression failed to converge to the exponential function for one cerebellar sham tDCS long preparation time dataset.

Stimulation (cerebellar anodal tDCS, cerebellar sham tDCS) x Preparation Time (short preparation time, long preparation time) x Rotation Direction (clockwise, counterclockwise) ANOVAs were run on rate constants. For all ANOVAs, when Mauchly's test of sphericity was significant, the Greenhouse-Geisser correction was used to adjust degrees of freedom. Partial η-squares were used to report ANOVA effect sizes, with values of 0.01 or less considered small, values between 0.01 and 0.09 considered medium; and values in excess of 0.25 considered large. Sidak corrections were used for post-hoc tests where necessary.

## Results

Figure 1 plots movement directions for all experimental blocks, collapsed across the long and the short preparation time conditions (Figure 1 top panel) and collapsed across the cerebellar anodal tDCS and the cerebellar sham tDCS conditions (Figure 1 bottom panel). Figure 2 plots movement directions for all experimental blocks separately for each preparation time condition (short preparation time condition shown in Figure 2 top panel, long preparation time condition shown in Figure 2 bottom panel).

**Figgue 1.**
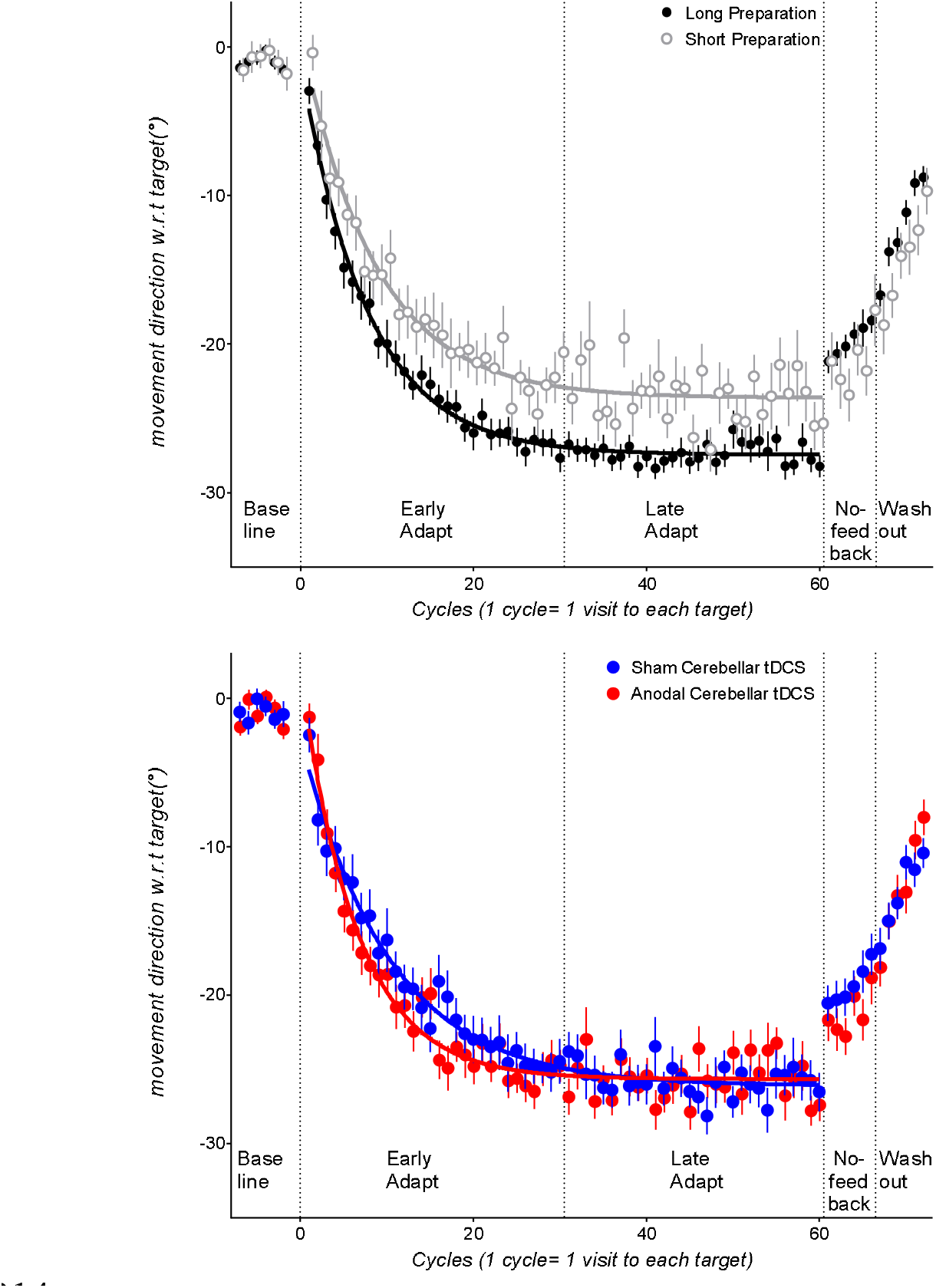
Top panel: cycle by cycle movement directions relative to the target, averaged across the short preparation time groups (clear circles), and the long preparation time groups (grey circles) (i.e., pooled across the cerebellar tDCS anodal and sh conditions). Bottom panel: cycle by cycle movement direction relative to the target, averaged across all participant groups w eceived cerebellar sham tDCS and cerebellar anodal tDCS (i.e., pooled across long and short preparation time groups). D from the counterclockwise rotation groups were sign-transformed to allow statistical comparisons between clockwise and co erclockwise groups. In the adaptation block, values closer to -30° indicate more complete error compensation. In the no-feedback block, values closer to -30° indicate more implicit remapping, as participants were instructed that the rotation was removed, and that they were to aim straight towards the target without visual feedback of their movement. Error bars indicate standard errors of the mean. Lines indicate group mean data fit to the single-rate exponential function for the adaptation phase.

**Figure 2.**
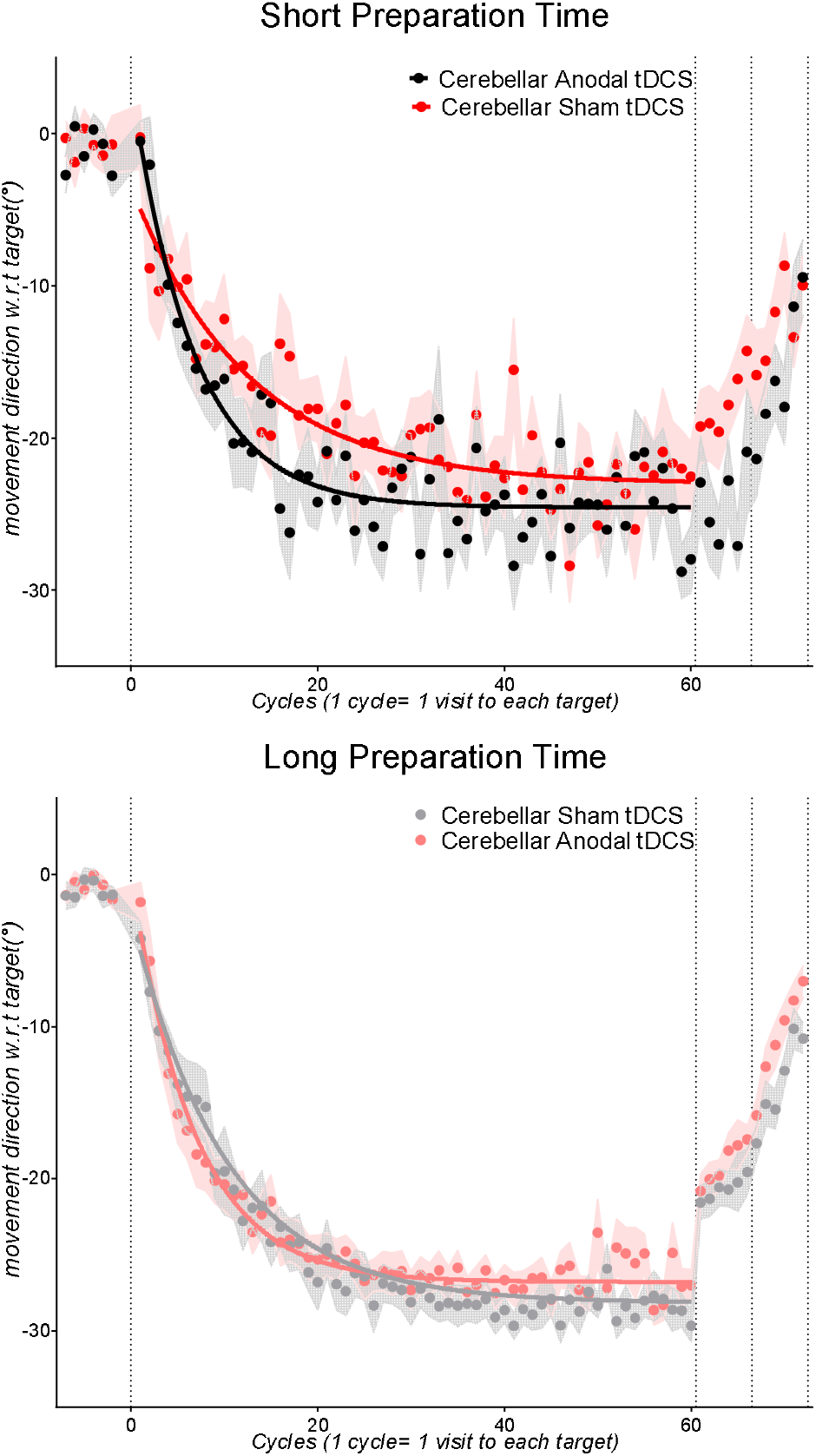
Top: Cycle by cycle movement directions with respect to the target for the short preparation time participants who re ed cerebellar anodal tDCS (red symbols) and cerebellar sham tDCS (black symbols). Bottom: cycle by cycle movement dir ions with respect to the target for the long preparation time participants who received cerebellar anodal tDCS (pink circles) and cerebellar sham tDCS (grey circles). Solid lines indicate group mean data fit to the single-rate exponential function for e adaptation phase.

Before the rotation was imposed, participants completed 48 baseline trials (i.e., 6 cycles; 6 visits to each tar. Participants tended to show a clockwise bias in this baseline phase (see Figure 1 top and bottom pa ls). To evaluate whether participant groups differed in accuracy of movement direction before the rotation was imposed, we ran a cycle (baseline cycle4, baseline cycle 5, baseline cycle 6) x Preparation Time (Short, Long) x Stim (Sham, Stim) ANOVA. Importantly, groups receiving anodal or sham cerebellar tDCS did not differ reliably in directional accuracy, as there was no significant main effect of stimulation. Movement directions also did not differ reliably between the long and the short preparation time conditions (non-significant main effect of Preparation Time, no significant interactions with preparation time, all p>0.2).

**Early adaptation:** After the 30° rotation was imposed, participants compensated for the error imposed by the rotation by moving in the opposite direction to the rotation (see Figure 1, where more compensation for a 30° clockwise rotation would be indicated by movements closer to 30°: data from the counterclockwise rotation conditions were sign transformed to allow collapsing with data from the clockwise rotation condition). To evaluate the effect of the tDCS and preparation time manipulations on the early phase of error compensation, a Stimulation (cerebellar anodal tDCS, cerebellar sham tDCS) x Preparation Time (short preparation time, long preparation time) x Cycle (Adaptation Cycle 1…Cycle 30) ANOVA was run. Movement directions became progressively closer to the adapted movement direction with increasing cycles, as shown by a significant main effect of Cycles, F(12,721.6) = 69.09, p = 0, partial η-squared = 0.53. Constraining preparation time resulted in less error compensation (see Figure 1 top panel, where better compensation for the rotation is indicated by movements closer to -30°), as shown by a significant main effect of Preparation Time, F(1,60) = 6.71, p = 0.012, partial η-squared = 1.1. Hence, shortening preparation time resulted in less error compensation in the early phase of adaptation, corroborating our previous results which showed that shortening preparation time can provide a sufficient assay of implicit learning [25]. We previously showed that shortening preparation time in this way resulted in similar rates and extents of error compensation to estimates of implicit learning obtained by subtracting aiming directions[14].

Similar to previous research [19, 21, 40], error compensation for the visuomotor rotation tended to be faster for cerebellar anodal tDCS than for sham tDCS (see Figure 1 bottom panel), (significant Cycles x Stim interaction, F(12,721.6) = 2.18, p = 0.011, partial η-squared = 0.03). Note however that this was only a moderate effect size, appeared considerably weaker than that found in previous studies[19, 21, 40].

**Late adaptation:** Stimulation (cerebellar anodal tDCS, cerebellar sham tDCS) x Preparation Time (short preparation time, long preparation time) x Cycle ANOVA was run on the late phase (cycles 31…60) of the adaptation block. Similar to our previous results [25], restricting preparation time resulted in less error compensation in the late adaptation phase (see top panel Figure 1), as shown by a significant main effect of Preparation Time, F(1,60) = 12.02, p = 0.001, partial η-squared = 0.16. There was a significant Cycles x Stim interaction, F(13.8,829.4) = 2, p = 0.016, partial η-squared = 0.03.

### Rate of adaptation quantified by rate constants

To guard against the possibility that results from analyses of mean movement directions resulted from individual differences in intrinsic directional biases in reaching movements, we additionally quantified error compensation in terms of rate constants obtained from fitting adaptation phase single subject data to a single-rate exponential model [19]. Preparation Time x Stimulation ANOVAs on rate constants showed a marginal main effect of stimulation, F(1,59) = 3.09, p = 0.084, partial η-squared = 0.05. This reflected a trend for larger mean rate constants (i.e., faster adaptation) with cerebellar anodal tDCS than with sham (see Figure 3A). Rate constants for the groups receiving anodal tDCS tended to be larger than rate constants for the groups receiving sham tDCS (see Figure 3A). The main effect of preparation time was not reliable. There were no other reliable main effects or interactions.

**Figure 3.**
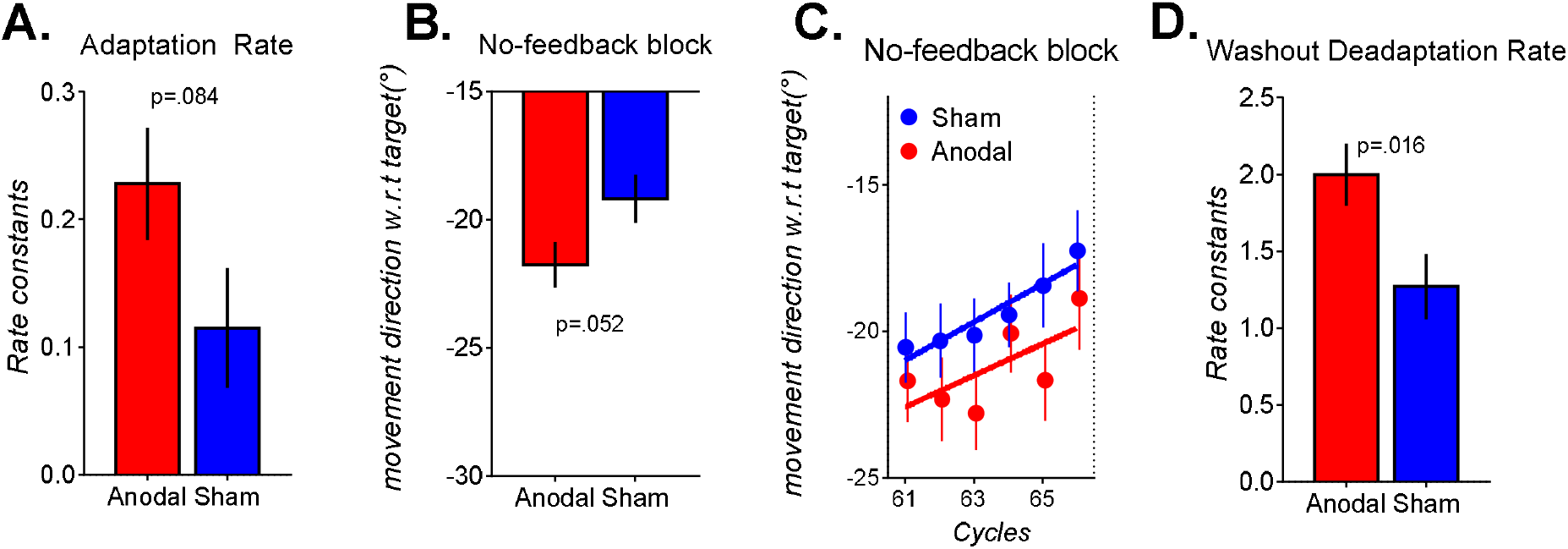
Group mean (err bars are SEM) for data f m the cerebellar anodal t CS condition (Stim, red and the cerebellar sha tDCS condition (Sham, blue). **A**: rate constants from fitting a single-rate exponential function to cycle-averaged movement directions from the adaptatio block—larger values indicate faster adaptation he rotation. **B & C.** Gro mean movement direction from the no-feedback block, averaged across cycles (B) or cycle-by-cycle (C)—values closer to -30° represent more adapted movements. **D**. Slopes from fitting a straight line to individual cycle-averaged movement directions in the washout block-larger values indicate faster de-adaptation to the unadapted state.

### Cerebellar anodal tDC increased implicit re apping

After exposure to the pertu ation in the adaptation b ck, participants were explicitly told the rotation was removed, and they sho d aim straight to the ta ts in the subsequent post-adaptation no-feedback block, following protocols from Taylor, Krakauer (1 These instructions were crucial to properly quantify *implicit* acquisition of sensorimotor maps resulting from adaptation to the rotation (thereafter termed *implicit remapping*), as movements that remained adapted despite explicit instructions that the rotation was no longer present are likely to reflect residual implicit learning after removing the influence of explicit learning. Explicit knowledge that the rotation has been removed results in an abrupt drop-off of adaptation from the last adaptation cycle to the first no-feedback cycle, as previously documented [25].

A Preparation Time (Long, Short) x Stimulation (anodal, sham) x Cycle (No Feedback Cycle 1… Cycle 6) ANOVA showed a significant main effect of Cycles, F(3.3,201.2) = 3.48, p = 0.013, partial η-squared = 0.05, as movements decayed slowly across cycles in the absence of visual feedback, corroborating previous results [14, 21, 41]. There was a marginal main effect of Stim, F(1,60) = 3.94, p = 0.052, partial η-squared = 0.06 (moderate effect size), as movements were overall more adapted with anodal tDCS than with sham (see Figure 3B, cycle averaged movement directions across the no-feedback block: anodal tDCS: -21.7°/-0.9°, sham tDCS: -19.1°/-0.9°). There was a significant Preparation Time x Stim interaction, F(1,60) = 10.39, p = 0.002, partial η-squared = 0.14. Follow-up ANOVAs were run separately for the Short and the Long preparation time datasets. For short preparation time, there was a significant main effect of stimulation, F(1,25) = 7.65, p = 0.01, partial η-squared = 0.23, as movements were more adapted in this no-feedback block with anodal stimulation than with sham stimulation (group mean of no-feedback cycles: sham tDCS =17.7°/-1.8°, anodal tDCS= 24.5°/-1.7°, p = .010, sidak-corrected, d=1.04, large effect size). For the long preparation time group, the main effect of stimulation and the cycle x stimulation interaction failed to reach significance.

**Washout:** Here, cursor feedback of movements was returned, and the error incurred by removal of the rotation became visible to participants. Participants thus rapidly returned movements to the un-adapted state (see Figure 1 washout phase). Preparation Time (Short, Long) x Stim (Anodal, Sham) x Cycles (Washout 1…Washout 6) ANOVA showed a significant Preparation Time x Stim interaction, F(1,60) = 6.85, p = 0.011, partial η-squared = 0.1 a significant Cycles x Stim interaction, F(4,245.6) = 2.4, p = 0.049, partial η-squared = 0.03, and a significant Cycles x Preparation Time x Stim interaction, F(4,245.6) = 2.76, p = 0.027, partial η-squared = 0.04. Follow-up ANOVAs were run separately for the short and long preparation time conditions. For the short Preparation Time groups, there was a significant Cycles x Stim interaction, F(4.4,114.9) = 2.49, p = 0.041, partial η-squared = 0.08, as participants in the anodal tDCS condition showed slower washout of adapted movements to the un-adapted state compared to sham tDCS. For the long preparation time condition, there was main effect of stimulation, F(1,35) = 5.61, p = 0.023, partial η-squared = 0.13, as movements in the anodal tDCS condition tended to be *less* adapted compared to sham (mean of washout cycles 1-6: sham tDCS: -13.6°/-0.9°, anodal tDCS -10.8°/-0.8°). These results corroborate that of Galea et al. (2011) who found a trend for faster washout with cerebellar anodal tDCS in their Experiment 1.

**Rate of deadaptation quantified by rate constants:** We fit individual cycle-averaged washout phase movement directions to a straight line to obtain slopes—this provides an assay of the rate of washout which is less influenced by differences in the intrinsic biases in reaching direction, as well as movement direction at the start of the washout phase. We ran a Preparation Time x Stim x Rotation Direction ANOVA on the slopes. There was a significant main effect of Stim F(1,60)=6.16, p =.016, partial η- squared = .09, as the rate of washout was faster with anodal tDCS than with sham (see Figure 3D).

## Discussion

The cerebellum has long been known to play a crucial role in adapting movements to perturbations of sensory feedback [7-9]. Previous work showed that increasing cerebellar excitability via non-invasive stimulation of the cerebellum speeds up error compensation during adaptation to perturbations such as rotated visual feedback [19, 21, 40], force-field perturbations of movement trajectories [22], as well as locomotor adaptation to split-belt treadmill walking [20]. However, to the best of our knowledge, none of these previous studies controlled for the use of explicit processes, such as volitionally applied compensatory strategies. Previous studies demonstrating the effects of cerebellar tDCS tended to be clearest in the initial stage of error compensation [21], which is now known to be dominated by explicit processes [14]. Given accumulating evidence for the cerebellum's role in predicting sensory events and fine-tuning of behavioral responses in many higher-order cognitive tasks [42], it was unclear whether previous findings of faster error compensation with cerebellar stimulation were due to the effects of cerebellar stimulation on explicit processes, implicit processes, or both. The current data show that despite reducing the amount of time available for movement preparation to suppress explicit strategy use, cerebellar anodal tDCS still increased the rate of adaptation. Furthermore, after perturbation removal and despite explicit knowledge that the rotation had already been removed, movements remained more adapted with cerebellar anodal tDCS than with sham tDCS. These results provide evidence of cerebellar involvement in implicit learning processes in adaptation to visuomotor rotations in healthy adults.

These current findings that increasing cerebellar excitability increased post-adaptation implicit remapping corroborates a large body of work in humans and non-human primates [3]. Previous studies have shown smaller aftereffects in patients with cerebellar degeneration [4, 7, 43, 44], however, these findings of smaller aftereffects might not reflect deficits in implicit error-based learning alone, as it is unclear whether patients and healthy controls had equivalent explicit knowledge of perturbation removal in those studies.

We also found that increasing cerebellar excitability with anodal tDCS increased the rate at which participants altered movements to (1) reduce errors resulting from a rotation in the adaptation phase, and (2) reduce errors resulting from sudden removal of a rotation after adapting movements to the rotation. This result corroborates that of previous work, although the effect of cerebellar anodal tDCS on error compensation here was substantially weaker in comparison to previous studies which did not control movement preparation time [45]. This might be due to differences in our experimental procedure, for example the stimulation intensity employed here (1.5mA) was weaker than that of previous studies (2mA) [18, 19, 21, 40]. Note however that a recent study that used a 2.0mA stimulation intensity found non-significant effect of cerebellar tDCS on sensorimotor adaptation [46], and recent evidence suggests that anodal tDCS intensities of 1.0 mA, 1.5mA and 2.0mA cause in similar effects on cortical excitability, at least in the motor cortex [47]. However, this comprehensive review by Jamil et al. (2016) shows that there are typically large individual differences in stimulation sensitivity, which might have contributed to the weak effect of cerebellar tDCS on error compensation in our results.

### Possible cerebellar involvement in explicit and implicit learning

Explicit and implicit processes are thought to work in tandem to compensate for errors resulting from perturbed sensory feedback [48-51]. It is possible that previously reported improvements in error compensation with cerebellar anodal tDCS might be partly driven by augmentation of explicit processes that result in the rapid error compensation early in sensorimotor adaptation [14]. The proposal that cerebellar anodal tDCS might alter explicit processes is consistent with reports of faster error compensation with cerebellar anodal tDCS in older adults [19], who have been widely documented to show slower error compensation as a result of poorer explicit learning [34, 52-55]. Poorer cerebellar function in older adults has also been linked to poorer explicit learning [56, 57]. There is evidence supporting cerebellar involvement in explicit learning: although cerebellar degeneration patients can employ an strategy when explicitly instructed to do so, they appear unable to spontaneously generate an explicit strategy [16], unlike healthy controls. Explicit learning is likely to be sensitive to reinforcement-based processes that influence movement selection[58]. The employment of reinforcement-based explicit processes is affected by cerebellar function, as recent studies show that although cerebellar degeneration patients can sometimes show residual ability to use compensatory mechanisms (e.g., online feedback and/or reinforcement mechanisms) to adapt movements to perturbed feedback that has been imposed gradually [59-63], they are poorer at learning from reinforcement [64, 65], possibly because of increased motor noise. The possibility that explicit processes are supported by cerebellar function is consistent with a growing body of work demonstrating the role of the cerebellum in predicting sensory events and fine-tuning of behavioral responses in many “non-motor” cognitive processes (for reviews, see [66, 67]. Thus, it seems likely that the cerebellum contributes to both explicit and implicit processes in sensorimotor adaptation[68].

There is evidence supporting the suggestion that distinct regions of the cerebellum support explicit and implicit processes in sensorimotor adaptation. For example, patients with posterior cerebellar lesions show deficits in the early part of error compensation thought to be primarily driven by explicit strategic processes, but not deficits in aftereffects thought to be driven by implicit processes [11]. In contrast, patients with superior cerebellar lesions showed more severe deficits in both the rate and extent of error compensation, as well as aftereffects, suggesting involvement of the superior cerebellum in implicit processes in sensorimotor adaptation [11]. Neuroimaging work in prism adaptation [69-71] also support the idea that distinct sub-regions of the cerebellum sub-serve implicit and explicit processes. In prism adaptation, the early phase of error compensation dominated by explicit processes is thought to be sub-served by a network encompassing the ventro-caudal dentate nucleus to the posterior parietal cortex [72, 73]. This proposal is supported by neuroimaging evidence showing greater activation of the ventro-caudal dentate nucleus and the posterior cortex of the cerebellum in the early phase of error compensation than in the late phase of error compensation [74]. Implicit processes are associated with greater cerebellar activation in the right anterior lobules IV/V in prism adaptation [75] and in lobule V and VI in adaptation to visuomotor rotations [76]. Our current findings do not allow us to speculate on which area of the cerebellum is affected by our cerebellar stimulation protocol, as tDCS effects are not focal. Employing concurrent cerebellar tDCS with fMRI whilst experimentally manipulating the use of explicit strategies might help illuminate how the cerebellum contributes to explicit processes during sensorimotor adaptation.

## Summary

Previous work using non-invasive brain stimulation demonstrated that the cerebellum plays a role in sensorimotor adaptation, however, because these studies did not dissociate explicit and implicit processes that occur during adaptation, it was unclear whether the cerebellum plays a role in implicit or explicit processes, or both. Here, we show that when explicit re-aiming processes is suppressed, increasing cerebellar excitability via anodal tDCS increases implicit remapping after adaptation to a 30° rotation. Thus, the cerebellum contributes to implicit sensorimotor remapping when people learn to compensate a visuomotor rotation.

## References

1. Wolpert DM, Miall RC, Kawato M. Internal models in the cerebellum. Trends Cogn Sci. 1998;2(9):338–47. PubMed PMID: 21227230.

2. Popa LS, Streng ML, Hewitt AL, Ebner TJ. The Errors of Our Ways: Understanding Error Representations in Cerebellar-Dependent Motor Learning. Cerebellum. 2016;15(2):93–103. DOI:10.1007/s12311-015-0685-5..

3. Brooks JX, Carriot J, Cullen KE. Learning to expect the unexpected: rapid updating in primate cerebellum during voluntary self-motion. Nat Neurosci. 2015.

4. Tseng YW, Diedrichsen J, Krakauer JW, Shadmehr R, Bastian AJ. Sensory prediction errors drive cerebellum-dependent adaptation of reaching. J Neurophysiol. 2007;98(1):54–62. DOI:10.1152/jn.00266.2007. PubMed PMID: 17507504.

5. Smith MA, Shadmehr R. Intact ability to learn internal models of arm dynamics in Huntington's disease but not cerebellar degeneration. Journal of Neurophysiology. 2005;93(5):2809–21. PubMed PMID: ISI:000228575200040.

6. Maschke M, Gomez CM, Ebner TJ, Konczak J. Hereditary cerebellar ataxia progressively impairs force adaptation during goal-directed arm movements. J Neurophysiol. 2004;91(1):230–8. DOI:10.1152/jn.00557.2003. PubMed PMID: 13679403.

7. Martin TA, Keating JG, Goodkin HP, Bastian AJ, Thach WT. Throwing while looking through prisms I. Focal olivocerebellar lesions impair adaptation. Brain. 1996;119(4):1183–98.

8. Weiner MJ, Hallett M, Funkenstein HH. Adaptation to lateral displacement of vision in patients with lesions of the central nervous system. Neurology. 1983;33(6):766–72.

9. Pisella L, Rossetti Y, Michel C, Rode G, Boisson D, Pélisson D, et al. Ipsidirectional impairment of prism adaptation after unilateral lesion of anterior cerebellum. Neurology. 2005;65(1):150–2. DOI:10.1212/01.wnl.0000167945.34177.5e.

10. Schlerf JE, Xu J, Klemfuss NM, Griffiths TL, Ivry RB. Individuals with cerebellar degeneration show similar adaptation deficits with large and small visuomotor errors. J Neurophysiol. 2013;109(4):1164–73. DOI:10.1152/jn.00654.2011. PubMed PMID: 23197450; PubMed Central PMCID: PMC3569142.

11. Werner S, Bock O, Gizewski ER, Schoch B, Timmann D. Visuomotor adaptive improvement and aftereffects are impaired differentially following cerebellar lesions in SCA and PICA territory. Experimental brain research. 2010;201(3):429–39. DOI:10.1007/s00221-009- 2052-6. PubMed PMID: 19885654; PubMed Central PMCID: PMC2832877.

12. Morton SM, Bastian AJ. Cerebellar contributions to locomotor adaptations during splitbelt treadmill walking. Journal of Neuroscience. 2006;26(36):9107–16.

13. Morton SM, Bastian AJ. Prism adaptation during walking generalizes to reaching and requires the cerebellum. Journal of Neurophysiology. 2004;92(4):2497–509. DOI:10.1152/jn.00129.2004.

14. Taylor JA, Krakauer JW, Ivry RB. Explicit and implicit contributions to learning in a sensorimotor adaptation task. The Journal of neuroscience: the official journal of the Society for Neuroscience. 2014;34(8):3023–32. DOI:10.1523/JNEUROSCI.3619-13.2014. PubMed PMID: 24553942; PubMed Central PMCID: PMC3931506.

15. Redding GM, Rossetti Y, Wallace B. Applications of prism adaptation: A tutorial in theory and method. Neuroscience and biobehavioral reviews. 2005;29(3):431–44.

16. Taylor JA, Klemfuss NM, Ivry RB. An Explicit Strategy Prevails When the Cerebellum Fails to Compute Movement Errors. Cerebellum. 2010;9(4):580–6. DOI:10.1007/s12311-010- 0201-x. PubMed PMID: WOS:000284955800012.

17. Bond KM, Taylor JA. Flexible explicit but rigid implicit learning in a visuomotor adaptation task. J Neurophysiol. 2015;113(10):3836–49. DOI:10.1152/jn.00009.2015. PubMed PMID: 25855690; PubMed Central PMCID: PMC4473515.

18. Yavari F, Mahdavi S, Towhidkhah F, Ahmadi-Pajouh MA, Ekhtiari H, Darainy M. Cerebellum as a forward but not inverse model in visuomotor adaptation task: a tDCS-based and modeling study. Experimental brain research. 2015. DOI:10.1007/s00221-015-4523-2. PubMed PMID: 26706039.

19. Hardwick RM, Celnik PA. Cerebellar direct current stimulation enhances motor learning in older adults. Neurobiol Aging. 2014;35(10):2217–21. DOI:10.1016/j.neurobiolaging.2014.03.030. PubMed PMID: 24792908; PubMed Central PMCID: PMC4087063.

20. Jayaram G, Tang B, Pallegadda R, Vasudevan EV, Celnik P, Bastian A. Modulating locomotor adaptation with cerebellar stimulation. J Neurophysiol. 2012;107(11):2950–7. DOI:10.1152/jn.00645.2011. PubMed PMID: 22378177; PubMed Central PMCID: PMC3378372.

21. Galea JM, Vazquez A, Pasricha N, de Xivry JJ, Celnik P. Dissociating the roles of the cerebellum and motor cortex during adaptive learning: the motor cortex retains what the cerebellum learns. Cereb Cortex. 2011;21(8):1761–70. DOI:10.1093/cercor/bhq246. PubMed PMID: 21139077; PubMed Central PMCID: PMC3138512.

22. Herzfeld DJ, Pastor D, Haith AM, Rossetti Y, Shadmehr R, O'Shea J. Contributions of the cerebellum and the motor cortex to acquisition and retention of motor memories. Neuroimage. 2014;98:147–58. DOI:10.1016/j.neuroimage.2014.04.076. PubMed PMID: 24816533; PubMed Central PMCID: PMC4099269.

23. Fernandez-Ruiz J, Wong W, Armstrong IT, Flanagan JR. Relation between reaction time and reach errors during visuomotor adaptation. Behav Brain Res. 2011;219(1):8–14. DOI:10.1016/j.bbr.2010.11.060. PubMed PMID: WOS:000289664100002.

24. Haith AM, Huberdeau DM, Krakauer JW. The influence of movement preparation time on the expression of visuomotor learning and savings. The Journal of neuroscience: the official journal of the Society for Neuroscience. 2015;35(13):5109–17. DOI:10.1523/JNEUROSCI.3869- 14.2015. PubMed PMID: 25834038.

25. Leow L-A, Marinovic W, Gunn R, Carroll TJ. Estimating the implicit component of visuomotor rotation learning by constraining movement preparation time. bioRxiv. 2016:082420.

26. Benson BL, Anguera JA, Seidler RD. A spatial explicit strategy reduces error but interferes with sensorimotor adaptation. J Neurophysiol. 2011;105(6):2843–51. DOI:10.1152/jn.00002.2011. PubMed PMID: 21451054; PubMed Central PMCID: PMC3118744.

27. Saijo N, Gomi H. Multiple motor learning strategies in visuomotor rotation. PLoS One. 2010;5(2):e9399. DOI:10.1371/journal.pone.0009399. PubMed PMID: 20195373; PubMed Central PMCID: PMC2827554.

28. Galea JM, Jayaram G, Ajagbe L, Celnik P. Modulation of Cerebellar Excitability by Polarity-Specific Noninvasive Direct Current Stimulation. 2009;29(28):9115–22.

29. Howard IS, Ingram JN, Wolpert DM. A modular planar robotic manipulandum with end-point torque control. Journal of Neuroscience Methods. 2009;181(2):199–211. DOI: http://dx.doi.org/10.1016/j.jneumeth.2009.05.005.

30. Schouten JF, Bekker JAM. Reaction time and accuracy. Acta Psychologica. 1967;27:143- 53. DOI:http://dx.doi.org/10.1016/0001-6918(67)90054-6.

31. Marinovic W, Tresilian JR, de Rugy A, Sidhu S, Riek S. Corticospinal modulation induced by sounds depends on action preparedness. The Journal of physiology. 2014;592(1):153- 69.

32. Haith AM, Pakpoor J, Krakauer JW. Independence of movement preparation and movement initiation. Journal of Neuroscience. 2016;36(10):3007–15. DOI:10.1523/JNEUROSCI.3245-15.2016.

33. Marinovic W, Plooy A, Tresilian JR. The time course of amplitude specification in brief interceptive actions. Exp Brain Res. 2008;188(2):275–88.

34. Heuer H, Hegele M. Adaptation to visuomotor rotations in younger and older adults. Psychol Aging. 2008;23(1):190–202. DOI:10.1037/0882-7974.23.1.190. PubMed PMID: 18361666.

35. Howard IS, Ingram JN, Franklin DW, Wolpert DM. Gone in 0.6 seconds: The encoding of motor memories depends on recent sensorimotor states. Journal of Neuroscience. 2012;32(37):12756–68. DOI:10.1523/JNEUROSCI.5909-11.2012.

36. Morehead JR, Ivry R. Intrinsic biases systematically affect visuomotor adaptation experiments. Neural Control of Movement; Charleston 2015.

37. Vindras P, Desmurget M, Viviani P, Error PV. Error Parsing in Visuomotor Pointing Reveals Independent Processing of Amplitude and Direction. 2008:1212–24.

38. Gordon J, Ghilardi MF, Ghez C. Accuracy of planar reaching movements - I. Independence of direction and extent variability. Exp Brain Res. 1994;99(1):97–111.

39. Zarahn E, Weston GD, Liang J, Mazzoni P, Krakauer JW. Explaining savings for visuomotor adaptation: linear time-invariant state-space models are not sufficient. J Neurophysiol. 2008;100(5):2537–48. DOI:10.1152/jn.90529.2008. PubMed PMID: 18596178; PubMed Central PMCID: PMC2585408.

40. Block H, Celnik P. Stimulating the cerebellum affects visuomotor adaptation but not intermanual transfer of learning. Cerebellum. 2013;12(6):781–93. DOI:10.1007/s12311-013-0486-7.

41. Kitago T, Ryan SL, Mazzoni P, Krakauer JW, Haith AM. Unlearning versus savings in visuomotor adaptation: Comparing effects of washout, passage of time and removal of errors on motor memory. Frontiers in Human Neuroscience. 2013;(JUN). DOI:10.3389/fnhum.2013.00307.

42. Peterburs J, Desmond JE. The role of the human cerebellum in performance monitoring. Curr Opin Neurobiol. 2016;40:38–44. DOI:http://dx.doi.org/10.1016/j.conb.2016.06.011.

43. Rabe K, Livne O, Gizewski ER, Aurich V, Beck A, Timmann D, et al. Adaptation to Visuomotor Rotation and Force Field Perturbation Is Correlated to Different Brain Areas in Patients With Cerebellar Degeneration. Journal of Neurophysiology. 2009;101(4):1961–71. DOI:10.1152/jn.91069.2008. PubMed PMID: WOS:000264465000027.

44. Werner S, Bock O, Timmann D. The effect of cerebellar cortical degeneration on adaptive plasticity and movement control. Experimental brain research. 2009;193(2):189–96. DOI:10.1007/s00221-008-1607-2. PubMed PMID: 18949468.

45. Schlerf JE, Galea JM, Spampinato D, Celnik PA. Laterality Differences in Cerebellar-Motor Cortex Connectivity. Cerebral Cortex. 2015;25(7):1827–34. DOI:10.1093/cercor/bht422.

46. Mamlins A, Hulst T, Donchin O, Timmann D, Claa²en J. EP 72. Cerebellar tDCS effects on the adaptation of arm reaching movements to force-field perturbations. Clinical Neurophysiology. 127(9):e269. DOI:10.1016/j.clinph.2016.05.123.

47. Jamil A, Batsikadze G, Kuo HI, Labruna L, Hasan A, Paulus W, et al. Systematic evaluation of the impact of stimulation intensity on neuroplastic after?effects induced by transcranial direct current stimulation. The Journal of Physiology. 2016.

48. Cunningham HA. Aiming error under transformed spatial mappings suggests a structure for visual-motor maps. J Exp Psychol Hum Percept Perform. 1989;15(3):493–506. PubMed PMID: 2527958.

49. Jakobson LS, Goodale MA. Trajectories of reaches to prismatically-displaced targets: evidence for "automatic" visuomotor recalibration. Experimental brain research. 1989;78(3):575- 87. DOI:10.1007/BF00230245. PubMed PMID: 2612600.

50. Uhlarik JJ. Role of cognitive factors on adaptation to prismatic displacement. J Exp Psychol. 1973;98(2):223–32. DOI:10.1037/h0034364. PubMed PMID: 4705623.

51. Redding GM, Wallace B. Cognitive interference in prism adaptation. Perception and Psychophysics. 1985;37(3):225–30.

52. Bruijn SM, Van Impe A, Duysens J, Swinnen SP. Split-belt walking: Adaptation differences between young and older adults. Journal of Neurophysiology. 2012;108(4):1149–57. DOI:10.1152/jn.00018.2012.

53. Cressman EK, Salomonczyk D, Henriques DY. Visuomotor adaptation and proprioceptive recalibration in older adults. Experimental brain research. 2010;205(4):533–44. DOI:10.1007/s00221-010-2392-2. PubMed PMID: 20717800.

54. Heuer H, Hegele M. Adaptation to direction-dependent visuo-motor rotations and its decay in younger and older adults. Acta psychologica. 2008;127(2):369–81. DOI:10.1016/j.actpsy.2007.07.006. PubMed PMID: 17826726.

55. Bock O, Schneider S. Sensorimotor adaptation in young and elderly humans. Neuroscience and biobehavioral reviews. 2002;26(7):761–7. PubMed PMID: 12470687.

56. Bernard JA, Seidler RD. Moving forward: Age effects on the cerebellum underlie cognitive and motor declines. Neuroscience and biobehavioral reviews. 2014;42:193–207. DOI:10.1016/j.neubiorev.2014.02.011.

57. Bernard JA, Seidler RD. Cerebellar contributions to visuomotor adaptation and motor sequence learning: An ALE meta-analysis. Frontiers in Human Neuroscience. 2013;(FEB). DOI:10.3389/fnhum.2013.00027.

58. Izawa J, Shadmehr R. Learning from sensory and reward prediction errors during motor adaptation. Plos Comput Biol. 2011;7(3):e1002012. DOI:10.1371/journal.pcbi.1002012. PubMed PMID: 21423711; PubMed Central PMCID: PMC3053313.

59. Hanajima R, Shadmehr R, Ohminami S, Tsutsumi R, Shirota Y, Shimizu T, et al. Modulation of error-sensitivity during a prism adaptation task in people with cerebellar degeneration. J Neurophysiol. 2015;114(4):2460–71. DOI:10.1152/jn.00145.2015. PubMed PMID: 26311179; PubMed Central PMCID: PMC4620141.

60. Criscimagna-Hemminger SE, Bastian AJ, Shadmehr R. Size of error affects cerebellar contributions to motor learning. J Neurophysiol. 2010;103(4):2275–84. DOI:10.1152/jn.00822.2009. PubMed PMID: 20164398; PubMed Central PMCID: PMC2853280.

61. Henriques DY, Filippopulos F, Straube A, Eggert T. The cerebellum is not necessary for visually driven recalibration of hand proprioception. Neuropsychologia. 2014;64C:195–204. DOI:10.1016/j.neuropsychologia.2014.09.029. PubMed PMID: 25278133.

62. Izawa J, Criscimagna-Hemminger SE, Shadmehr R. Cerebellar contributions to reach adaptation and learning sensory consequences of action. The Journal of neuroscience: the official journal of the Society for Neuroscience. 2012;32(12):4230–9. DOI:10.1523/JNEUROSCI.6353-11.2012. PubMed PMID: 22442085; PubMed Central PMCID: PMC3326584.

63. Gibo TL, Criscimagna-Hemminger SE, Okamura AM, Bastian AJ. Cerebellar motor learning: are environment dynamics more important than error size? Journal of Neurophysiology. 2013;110(2):322–33.

64. Therrien AS, Wolpert DM, Bastian AJ. Effective reinforcement learning following cerebellar damage requires a balance between exploration and motor noise. Brain. 2015:awv329.

65. McDougle SD, Boggess MJ, Crossley MJ, Parvin D, Ivry RB, Taylor JA. Credit assignment in movement-dependent reinforcement learning. Proceedings of the National Academy of Sciences of the United States of America. 2016;113(24):6797–802. DOI:10.1073/pnas.1523669113.

66. Koziol LF, Budding D, Andreasen N, D'Arrigo S, Bulgheroni S, Imamizu H, et al. Consensus paper: The cerebellum's role in movement and cognition. Cerebellum. 2014;13(1):151–77. DOI:10.1007/s12311-013-0511-x.

67. Stoodley CJ. The cerebellum and cognition: Evidence from functional imaging studies. Cerebellum. 2012;11(2):352–65. DOI:10.1007/s12311-011-0260-7.

68. Taylor JA, Ivry RB. Cerebellar and Prefrontal Cortex Contributions to Adaptation, Strategies, and Reinforcement Learning. Prog Brain Res2014. p. 217–53.

69. Luauté J, Schwartz S, Rossetti Y, Spiridon M, Rode G, Boisson D, et al. Dynamic changes in brain activity during prism adaptation. Journal of Neuroscience. 2009;29(1):169–78. DOI:10.1523/JNEUROSCI.3054-08.2009 10.1007/S00221-008-1303-2;. Suchan, B., Yágüez, L., Wunderlich, G., Canavan, A.G., Herzog, H., Tellmann, L., Hömberg, V., Seitz, R.J., Hemispheric dissociation of visual-pattern processing and visual rotation (2002) Behav Brain Res, 136, pp. 533–544; Weiner, M.J., Hallett, M., Funkenstein, H.H., Adaptation to lateral displacement of vision in patients with lesions of the central nervous system (1983) Neurology, 33, pp. 766–772; Zeffiro, T., Adaptation of visually-guided reaching to laterally displaced vision: A regional cerebral blood flow study [abstract] (1995) Hum Brain Mapp, (SUPPL. 1), p. 333.

70. Chapman HL, Eramudugolla R, Gavrilescu M, Strudwick MW, Loftus A, Cunnington R, et al. Neural mechanisms underlying spatial realignment during adaptation to optical wedge prisms. Neuropsychologia. 2010;48(9):2595–601.

71. Danckert J, Ferber S, Goodale MA. Direct effects of prismatic lenses on visuomotor control: An event-related functional MRI study. Eur J Neurosci. 2008;28(8):1696–704. DOI:10.1111/j.1460-9568.2008.06460.x.

72. Clower DM, Hoffman JM, Votaw JR, Faber TL, Woods RP, Alexander GE. Role of posterior parietal cortex in the recalibration of visually guided reaching. Nature. 1996;383(6601):618–21.

73. Clower DM, West Ra, Lynch JC, Strick PL. The inferior parietal lobule is the target of output from the superior colliculus, hippocampus, and cerebellum. JNeurosci. 2001;21(1529-2401 SB - IM):6283–91. DOI:21/16/6283 [pii].

74. Küper M, Wünnemann MJS, Thürling M, Stefanescu RM, Maderwald S, Elles HG, et al. Activation of the cerebellar cortex and the dentate nucleus in a prism adaptation fMRI study. Hum Brain Mapp. 2014;35(4):1574–86. DOI:10.1002/hbm.22274.

75. Luauté J, Michel C, Rode G, Pisella L, Jacquin-Courtois S, Costes N, et al. Functional anatomy of the therapeutic effects of prism adaptation on left neglect. Neurology. 2006;66(12):1859–67. DOI:10.1212/01.wnl.0000219614.33171.01.

76. Schlerf J, Ivry RB, Diedrichsen J. Encoding of sensory prediction errors in the human cerebellum. The Journal of neuroscience: the official journal of the Society for Neuroscience. 2012;32(14):4913–22. DOI:10.1523/JNEUROSCI.4504-11.2012. PubMed PMID: 22492047; PubMed Central PMCID: PMC4332713.

